# Microbial community-level features linked to divergent carbon flows during early litter decomposition in a constant environment

**DOI:** 10.1101/659383

**Authors:** Renee Johansen, Michaeline Albright, Deanna Lopez, La Verne Gallegos-Graves, Andreas Runde, Rebecca Mueller, Alex Washburne, Thomas Yoshida, John Dunbar

**Author notes:** **Corresponding author:** John Dunbar. Biosciences Division, Los Alamos National Laboratory, Mailstop M888, Los Alamos, NM 87545.

## Abstract

During plant litter decomposition in soils, carbon has two general fates: return to the atmosphere via microbial respiration or transport into soil where long-term storage may occur. Discovering microbial community features that drive carbon fate from litter decomposition may improve modeling and management of soil carbon. This concept assumes there are features (or underlying processes) that are widespread among disparate communities, and therefore amenable to modeling. We tested this assumption using an epidemiological approach in which two contrasting patterns of carbon flow in laboratory microcosms were delineated as functional states and diverse microbial communities representing each state were compared to discover shared features linked to carbon fate. Microbial communities from 206 soil samples from the southwestern United States were inoculated on plant litter in microcosms, and carbon flow was measured as cumulative carbon dioxide (CO_2_) and dissolved organic carbon (DOC) after 44 days. Carbon flow varied widely among the microcosms, with a 2-fold range in cumulative CO_2_ efflux and a 5-fold range in DOC quantity. Bacteria, not fungi, were the strongest drivers of DOC variation. The most significant community-level feature linked to DOC abundance was bacterial richness—the same feature linked to carbon fate in human-gut microbiome studies. This proof-of-principle study under controlled conditions suggests common features driving carbon flow in disparate microbial communities can be identified, motivating further exploration of underlying mechanisms that may influence carbon fate in natural ecosystems.

## Introduction

The decomposition of plant litter in soil is an ecosystem function with profound consequences. Litter decomposition recycles nutrients and directs photosynthetically fixed carbon into various carbon pools. Decomposition is primarily mediated by microorganisms. However in conventional models, physiochemical factors and plant traits are the primary controls over decomposition rates and soil carbon flow (1). An emerging view posits a stronger role for microbial composition (1–3) because microbial communities are not always functionally equivalent (4). If common features driving key functional outcomes occur in disparate microbial communities across diverse ecosystems, the mechanisms represented by the features could be incorporated into models to improve predictions of climate feedbacks and management strategies. We define “features” as taxa, gene families, metabolic pathways, or any other unit of interest measured by ‘omics data. Features are an entry point to discover mechanisms, which are the focus of modeling. Identifying robust community features that drive functional outcomes requires a large collection of communities that vary in composition and represent a substantial functional distribution from which contrasting functional states can be delineated. This approach is standard in epidemiology, where a large population is prescreened for subgroups that represent well-defined “healthy” versus “sick” functional states, for example.

Performing this type of study of variation in carbon flow patterns from litter decomposition in *natural* ecosystems is impeded by several logistical challenges. First, the geographic scale required to capture sufficient variation in community composition and function is unknown. Second, microbial composition is entangled with other ecosystem and microhabitat variables that obscure cause-effect relationships between microbial community features and carbon flow patterns. Third, obtaining functional measurements (e.g. release of CO_2_ or dissolved organic carbon) for a sufficient number of samples in the field is impractical.

A useful first step is to assess microbial-driven variation in decomposition outcomes in a simplified laboratory ecosystem (microcosms). This approach sacrifices some realism but enables tests of fundamental concepts and allows isolation of microbial community composition as the sole independent variable. With microcosms, hundreds of communities can be examined; the large scale reduces the risk of discovering eccentric features in a few communities that have little relevance at broader scales. Moreover, reliable measurements of carbon flow are feasible. The use of microcosms eliminates the spatial heterogeneity in edaphic factors that routinely confounds attempts to attribute carbon fate to microbial community features in field studies. Proof-of-principle studies with laboratory ecosystems can justify and guide more targeted validation studies in natural ecosystems (5, 6).

We measured carbon flow during the early phase of plant litter decomposition by quantifying dissolved organic carbon (DOC) and carbon dioxide (CO_2_) in microcosms under constant conditions. To obtain a large number of distinct source soil communities, we exploited the observation that microbial communities vary over large geographic scales (7). We collected 206 soil samples from 9 states in the Western U.S. encompassing 12 million km_2_ (Suppl. Fig. 1). We inoculated the communities into 618 (206 x 3 replicates) laboratory microcosms containing sterilized sand and plant litter (ponderosa pine needles, ground in a homogenous mixture). Plant litter decomposition is generally viewed as a two-stage process comprising an initial fast phase dominated by weedy microbial taxa, and a subsequent slow phase driven by taxa better equipped to decompose lignin and other recalcitrant substrates (8, 9). For an initial test of the concept of microbially driven variation in carbon-flow from litter, we focused on the early (fast) phase of plant litter decomposition, measuring cumulative carbon outputs after 44 days of decomposition. We then used the resulting distribution of DOC to delineate two functional states (“high” and “low” DOC) representing the most divergent levels of DOC for community profiling and feature analysis.

## Materials and methods

### Initial soil collection for microbial inoculum

Soil samples were collected from 206 locations throughout the southwestern United States between February and April, 2015 (Suppl. Fig. 1). The goal of this study was not to relate functional outcomes to detailed characteristics of the environments from which the soils were collected. Therefore, a randomized collection scheme was not used, as this would have substantially increased the cost and logistical burden of sample collection without benefit. Samples were typically collected at locations approximately 80 km apart, at least 15 meters from roadways, from the top 3cm of the soil surface after removal of surface litter (if any). Samples were collected in sterile 50-ml screw-cap tubes, and immediately stored on ice. The location of each sample was recorded by GPS and photographed to facilitate description of the major ecosystem types from which samples were obtained. GPS coordinates are available upon request.

### Microcosm construction and CO_2_ sampling

Microcosms were constructed using 125ml serum bottles. Each bottle contained approximately 5g of sand, and 0.12g of pine litter (dried pine needles), which had been ground in a Wiley Mill (Thomas Scientific, Swedesboro, NJ, USA). The microcosms were sterilized by autoclaving three times for 1 hour, with at least an 8-hour resting interval between each autoclave cycle. Microbial community inoculum was extracted from each soil sample (n=206) by suspending one gram of soil in 9ml of phosphate-buffered saline (PBS), then generating a 1000-fold dilution in PBS amended with NH_4_NO_3_ at 4.8mg/ml. Three microcosms per soil sample (n=618) each received 1.3 mls of inoculum, pipetted directly onto 0.02g of pine litter. Negative control microcosms, used to confirm the efficacy of sterilization, received the same quantities of PBS and NH_4_NO_3_, but no microbial communities. Sealed microcosms were incubated at 25°C in the dark for 14 days to equilibrate the communities. CO_2_ was evacuated using a vacuum pump on days 3 and 7, and replaced with sterile-filtered air. On day 14, a further 0.1 g of litter, which had been sterilized by three rounds of autoclaving, was added to each microcosm and microcosms were sealed with Teflon-lined crimp caps. The microcosms were incubated at 25°C in the dark for a further 30 days. During this time, CO_2_ was measured by gas chromatography using an Agilent Technologies 490 Micro GC (Santa Clara, CA, USA) on days 2, 5, 9, 16, 23 and 30. After each measurement, the headspace air was evacuated with a vacuum pump and replaced with sterile-filtered air. Cumulative CO_2_ was calculated by summing up the CO_2_ quantities recorded at each measurement point. CO_2_ data was used for only 289 microcosms, after it was discovered that the caps on the remaining microcosms were flawed and had been inadequately sealed.

### Dissolved organic carbon (DOC) and litter community sampling

After the 44-day (total) incubation, microcosms were destructively sampled to measure DOC and community composition. DOC extractions were performed using a rapid, gentle washing procedure to avoid measurement artifacts arising from microbial growth or microbial cell disruption. Specifically, 5ml of sterile deionized water was added to each microcosm, swirled manually for 30 seconds, then transferred by pipet to two 2-ml microfuge tubes. The tubes were centrifuged at 16,400xg for 4 minutes. The supernatants were combined and sterilized by filtration through a 0.2μm filter. The concentration of DOC in each sample was measured on an OI Analytical model 1010 wet oxidation TOC analyzer (Xylem Inc., Rye Brook, NJ, USA), calibrated daily. DOC data was obtained for 611 of 618 microcosms (7 samples were compromised during processing). Following DOC sampling, material (sand and litter) from each microcosm was frozen at −80°C for DNA extraction.

### Bacterial and fungal community taxonomic profiling

Samples for community profiling were down-selected based on the mean DOC quantity of each set of three replicates. Samples from replicate microcosms with the highest mean DOC (n=192, 64 soils x 3 replicates) and lowest mean DOC (n=192, 64 soils x 3) outputs were selected for DNA extraction and sequencing. These samples represent the two tails of the DOC distribution (Suppl. Fig. 3). DNA extractions were performed using a DNeasy PowerSoil 96-well plate DNA extraction kit (Qiagen, Hilden, Germany). The standard protocol was used with the following two exceptions: 1) 0.3 grams of material was used per extraction; 2) bead beating was conducted using a Spex Certiprep 2000 Geno/Grinder (Spex SamplePrep, Metuchen, NJ, USA) for three minutes at 1900 strokes/minute. DNA samples were quantified with an Invitrogen Quant-iT™ ds DNA Assay Kit (Thermo Fisher Scientific, Eugene, OR, USA) on a BioTek Synergy HI Hybrid Reader (Winooski, VT, USA). DNA quantities were used as a proxy for total microbial biomass. For comparisons of biomass between high and low DOC groups, and for correlation between biomass and DOC quantity, three samples greater than 3 standard deviations from the mean were removed as outliers, and one sample failed to yield DNA. This left 190 samples for each DOC group. PCR templates were prepared by diluting an aliquot of each DNA stock in sterile water to 1ng/μl. The bacterial (and archaeal) 16S rRNA gene (V3-V4 region) was amplified using primers 515f-R806 (10). Henceforth, archaeal sequences were analyzed with bacterial sequences. The fungal 28S rRNA gene (D2 hypervariable region) was amplified using the LR22R primer (11) and the reverse LR3 primer (12). Preparation for Illumina high-throughput sequencing was undertaken using a two-step approach, similar to that performed by Mueller et al. (13), with Phusion Hot Start II High Fidelity DNA polymerase (Thermo Fisher Scientific, Vilnius, Lithuania). In the first PCR, unique 6 bp barcodes were inserted into the forward and reverse primer in a combinatorial approach over 22 cycles with an annealing temperature of 60°C (14). The second PCR added Illumina-specific sequences over 10 cycles with an annealing temperature of 65°C. Amplicons were cleaned using a Mo bio UltraClean PCR clean-up kit (Carlsbad, CA, USA), quantified using the same procedure as for the extracted DNA, and then pooled at a concentration of 10ng each. The pooled samples were further cleaned and concentrated using the Mobio UltraClean PCR clean-up kit. All clean ups were undertaken as per the manufacturer’s instructions with the following modifications: binding buffer amount was reduced from 5X to 3X sample volume, and final elutions were performed in 50 μl Elution Buffer. A bioanalyzer was used to assess DNA quality, concentration was verified using qPCR, and paired-end 250 bp reads were obtained using an Illumina MiSeq sequencer at Los Alamos National Laboratory.

Bacterial and fungal sequences were merged with PEAR v 9.6 (15), quality filtered to remove sequences with 1% or more low-quality (q20) bases, and demultiplexed using QIIME (16) allowing no mismatches to the barcode or primer sequence. Further processing was undertaken with UPARSE (17). Sequences with an error rate greater than 0.5 were removed, remaining sequences were dereplicated, singletons were excluded from clustering, OTU clustering was performed at 97% and putative chimeras were identified *de novo* using UCHIME (18). Bacterial and fungal OTUs were classified using the Ribosomal Database Project (RDP) classifier (19). The OTUs that were not classified as bacteria or fungi with 100% confidence were removed from the dataset. Bacterial OTUs also had to have a phylum classification confidence level of at least 80% to remain in the dataset. Following quality control and classification, 9,576,525 sequences from 349 microcosms were obtained for bacteria and 13,124,107 sequences from 377 microcosms were obtained for fungi. These sorted into 2,527 OTUs for bacteria (an average of 275 per microcosm, SE = 8) and 753 OTUs for fungi (an average of 47 per microcosm, SE = 1). Sequence data has been deposited in the NCBI Sequence Read Archive (SRP151768). All other data including OTU tables are available upon request.

### Fungal:bacterial ratio

The bacterial content of the DNA extracted from microcosm material was determined using quantitative PCR (qPCR). For this exercise, a subset of random samples in the high and low DOC groups was chosen, with one replicate of each set of three randomly selected (*n* = 37 for low DOC communities, *n* = 35 for high DOC communities). We used primers EUB 338 (20) and EUB 518 (21) as described by Castro et al. (22) with the Biorad iQ SyBr Green Supermix on a BioRad CFX Connect Real-Time System (BioRad, Hercules, CA). DNA templates were normalized to 1.0 ng/μl and the annealing temperature was 55°C. Serial dilutions of genomic DNA from *Burkholderia thailandensis* E264 (ATCC 70038) were used to create a six-point calibration for qPCR DNA quantification. Melt curves were generated for every run to detect potential false positives. The 16S rRNA gene copy numbers determined from qPCR were converted to mass of bacterial DNA using an assumption of 4 copies (average) per genome, an average genome size of 3.5MB, and a mass conversion factor of 1.079375623^−9^ pg/bp. After computing the bacterial fraction of the total DNA, the remainder of the DNA was assumed to represent fungal DNA. This assumption was reasonable because the DNA obtained from the microcosms arose predominantly from the profuse fungal and bacterial mats that grew on the plant litter in every microcosm (attempts to extract DNA either from the original inoculum suspensions or from the sterile plant litter did not recover measurable quantities of DNA). The calculated masses of fungal DNA and bacterial DNA were used to compute the fungal:bacterial ratio. Three samples were removed as outliers as they were more than three standard deviations from the mean, leaving 34 samples in the high DOC group and 35 samples in the low DOC group.

### DOC binding assay

DOC binding potential was assessed for the DOC from one randomly-chosen replicate from all the high DOC (*n* = 64) and all the low DOC (*n* = 64) communities. For each sample, 0.5 ml of extracted DOC was added to 1ml of sterile water (creating a dilution factor of 3) and 0.3 grams of aluminum oxide (AlO_3_). Samples were gently mixed using a Thermolyne rocker (Barnstead/Thermolyne, Dubuque, IA, USA) at maximum tilt for 30 minutes and then centrifuged at maximum speed (16,100 xg) for 5 minutes. Supernatant was transferred to a new tube and stored at −20 C until DOC quantification on a TOC analyzer. The percentage of bound DOC was calculated as 100% x (DOC_post-binding_ x dilutionFactor)/DOC_pre-binding_.

### Statistical analyses

While initially each microbial inoculum was used in three replicate microcosms, the individual microcosms were used as independent samples in all statistical analyses. They were treated as individual samples because the initial inoculum replicates diverged substantially in community composition by the conclusion of the experiment. Additionally, compositional analyses were run on each set of replicates independently to confirm that our conclusions were robust irrespective of how replicates were treated.

Community composition analyses were performed with rarefied data unless otherwise stated. Bacterial communities were rarefied to 1023 sequences per sample, after which 311 samples remained for analysis. Fungal communities were rarefied to 2032 sequences, after which 345 samples remained for analysis. Bacterial (*n* = 311) and fungal (*n* = 345) richness and diversity (Shannon-Wiener index) were compared across high and low DOC groups using t-tests. Bray-Curtis dissimilarity matrices for bacterial and fungal communities were computed using log-transformed data for bacteria and for fungi in the R library vegan v 2.4-3 (23). Next, we used Pearson’s correlations to assess correspondence between community composition and DOC concentration for each of the 6 ordination dimensions. The strongest relationship was seen in the first (r = −0.46, *P* = < 0.001) and second (r = −0.44, *P* = <0.001) dimensions for bacteria. For fungi, the strongest relationship was seen in the fourth (r = 0.32, *P* = <0.001) and fifth (r = −0.21, *P* = <0.001) dimensions. We performed a permutational multivariate analysis of variance (PERMANOVA) (24) to assess whether the communities from high and low DOC groups differed (vegan v 2.4-3) (23). To quantify the relative variability within each DOC group (i.e. high and low), we measured the average distance to the centroid within each DOC group using a test for homogeneity of dispersion (24) (*vegan* v 2.4-3 package R) (23). This revealed that although dispersion was significantly different between high and low DOC microcosms for both bacteria (F_1,309_ = 69.56, *P* < 0.001) and fungi (F_1,343_ = 6, *P* < 0.05), the difference in dispersion between the high and low DOC groups was small. For fungi, the average distance to the centroid was 0.57 for the high DOC microcosms and 0.55 for the low. For bacteria, the average distance to the centroid was 0.51 for high and 0.46 for low DOC microcosms. The relationship between community composition and DOC quantity (instead of high vs low groupings) was also investigated using Mantel tests based on 999 permutations with log-transformed data. For bacteria, 308 samples were used, for fungi, 343 samples were used. Here, DOC values associated with each sample were used to generate a Euclidean distance matrix, used in Mantel tests with Bray-Curtis distance matrices for bacteria and fungi (*ecodist* package) (25).

To further compare community composition and DOC quantity, for high and low DOC groups, OTU sequences were grouped phylogenetically, for bacteria to family level and order level, and for fungi to order level. For fungal orders and bacterial families, OTUs were only used that could be phylogenetically assigned with at least a 70% confidence level from the RDP Classifier. For bacterial orders to be included, a classification confidence level of 90% was required. Family level comparisons were not made for fungi due to low classification confidence levels. Due to the large number of taxa present in the microcosms, we chose to perform further statistical analysis on only the most abundant taxa. For each bacterial and fungal order that comprised on average at least 0.1% of the sequences, and for each bacterial family that comprised on average at least 1% of the sequences of either (or both) the high or low DOC groups, we compared the proportion of sequences present in high versus low DOC groups using t-tests. For this analysis, the community composition matrices were not rarefied. For analysis of bacterial Orders, samples with less than 1063 sequences were removed, and for Families, samples with less than 1125 sequences were removed, in each case leaving 309 samples for this analysis. For fungi, samples with less than 1030 sequences were removed leaving 367 samples for this analysis.

Finally, we tested the strength of the relationship between various community features and DOC quantity using Pearson’s correlations. The features included fungal:bacterial ratios (n = 69), richness and Shannon diversity for bacteria (n = 308) and fungi (n = 343), total final biomass for fungi and for bacteria (n = 69), and total final biomass for fungi and bacteria combined (n = 377). All statistical analyses were performed using R v 3.3.3 (26).

## Results

### Variation in carbon quantity and quality

The quantity of CO_2_ produced by the microbial communities varied approximately 2-fold, between 160 and 345 mg/g of litter (Fig.1A). DOC accumulation showed a greater range, with between 3 and 18 mg/g of litter measured upon microcosm destruction (Fig.1B). The CO_2_ and DOC outputs from decomposition were negatively correlated (R_2_ = 0.16, *P* = <0.001; Suppl. Fig. 2). The DOC distribution was used to delineate two contrasting functional states. High and low DOC sample groups were delineated, respectively, as the tails of the DOC distribution (Suppl. Fig. 3) and were balanced by requiring each group to contain 192 samples (i.e., all 3 replicate communities derived from 64 source soils). The high and low DOC groups varied not only in DOC abundance but also in DOC composition, as indicated by a mineral-binding assay. The fraction of DOC binding to aluminum oxide ranged from 16.9% - 55.8% among the subset of DOC samples tested. Communities with high quantities of DOC had, on average, DOC with significantly greater potential for mineral-binding (two-tailed *t*-test, t_122_ = 2.8, *P* = 0.006; Fig. 2).

**Fig. 1.**
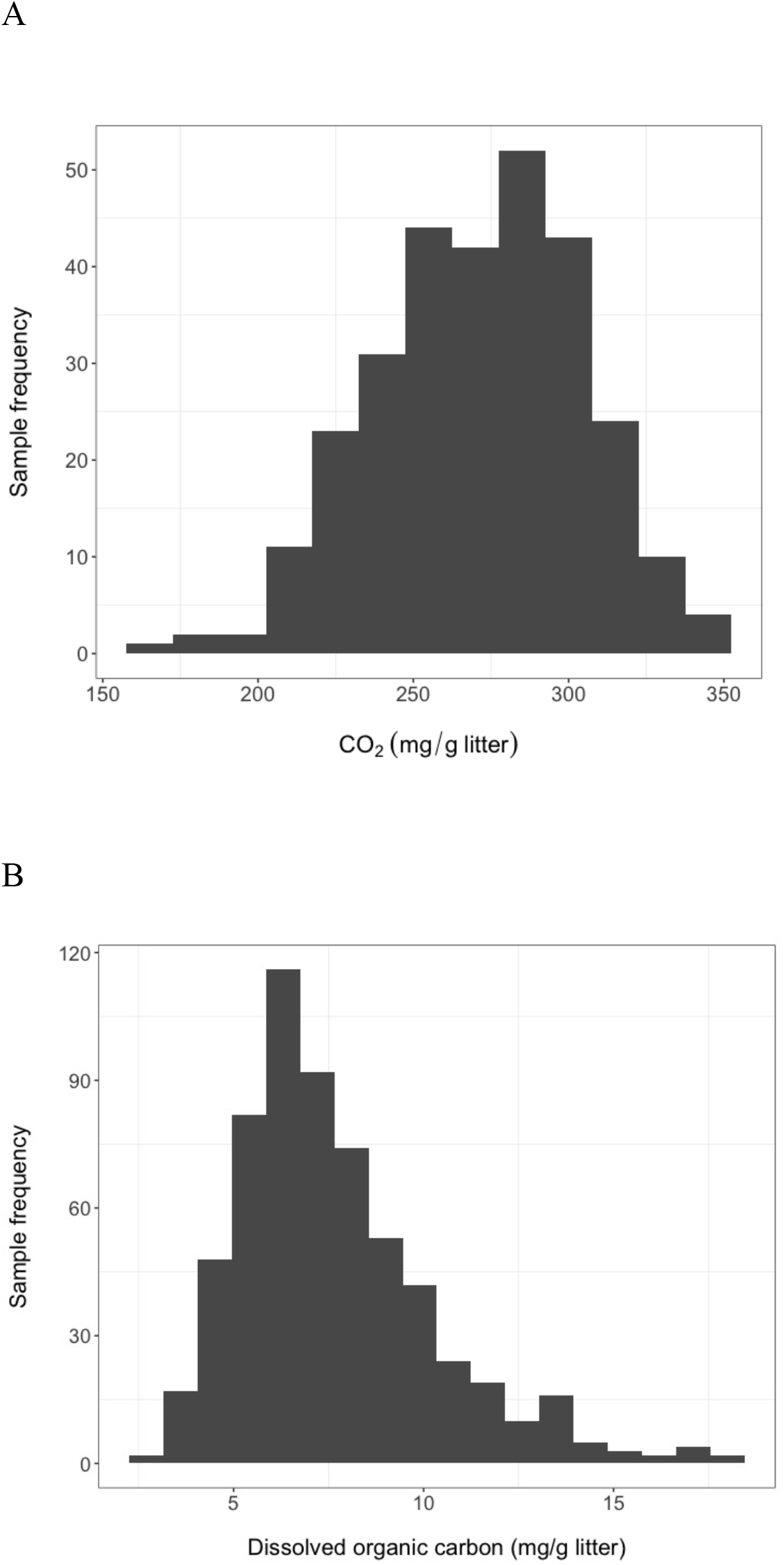
Carbon dioxide (*n* = 289) (A) and dissolved organic carbon (*n* = 611) (B) abundance from different microbial communities during 44 days of pine litter decomposition in microcosms

**Fig. 2.**
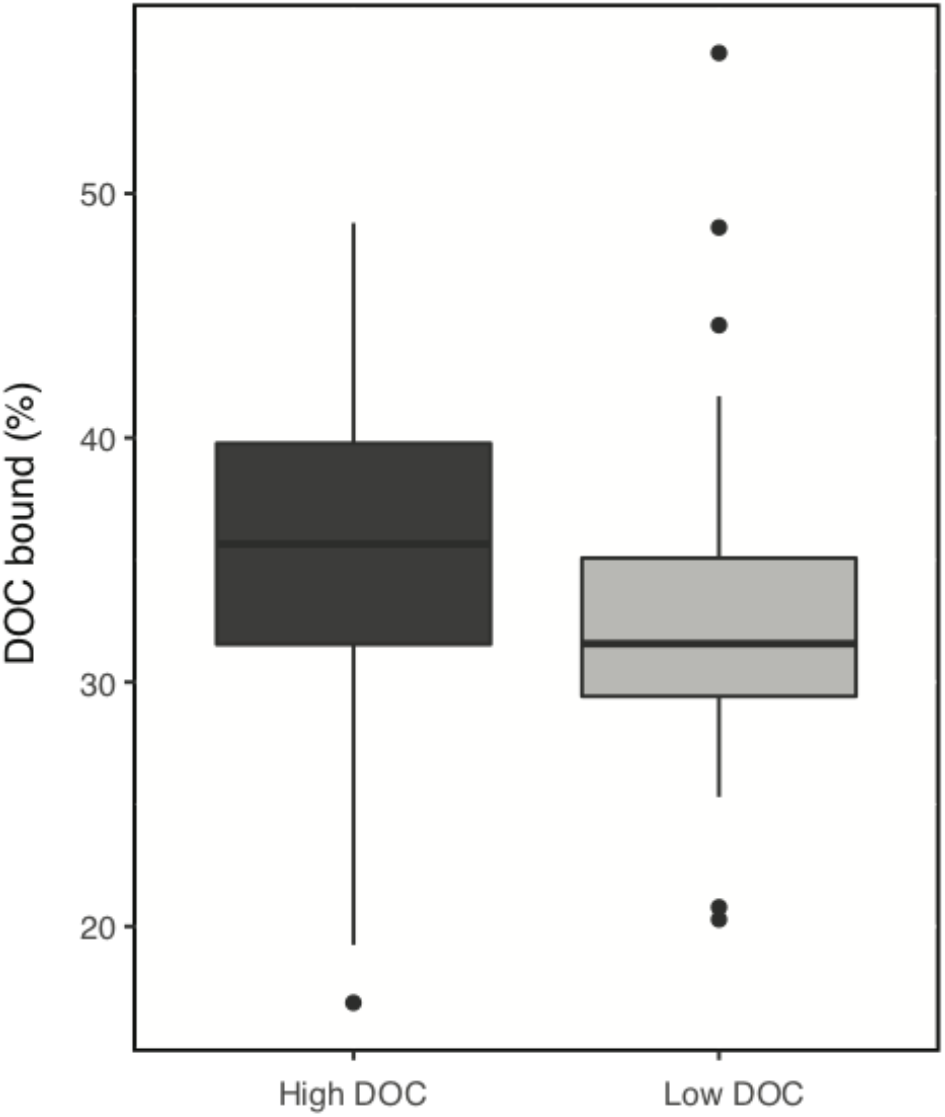
Proportion of dissolved organic carbon (DOC) that binds to aluminum oxide from high and low DOC microcosms. A greater proportion of DOC binds from high DOC than low DOC microcosms (*P* = 0.006, *n* = 128). DOC was produced by microbial communities during 44 days of decomposition of pine litter in microcosms.

### Geographic location of source soils yielding high and low DOC in microcosms

Source soil samples yielding high versus low DOC communities in our study were geographically intermingled (Suppl. Fig. 1) and co-occurred less than 30m apart at 14% of 49 geographic locations where two or more soil samples were collected from the same site. The source soils were obtained from eight ecosystem types defined broadly by dominant and minor plant types or by agricultural land-use (Table 1). The ecosystem type from which source soils were obtained significantly influenced the probability of observing high or low DOC abundance (ANOVA, F=28.87, P<0.0001) in the microcosm experiment. However, both functional outcomes occurred among source soils within each of five ecosystem types (Table 1), fulfilling the objective of acquiring contrasting functional states from multiple ecosystem types to search for common microbial community features.

**Table 1.** Number of samples from each ecosystem type which placed in medium, high and low DOC categories.

| **Ecosystem type<sup>a</sup>** | **Samples per DOC category** |  |  |
| --- | --- | --- | --- |
|  | **Medium** | **High** | **Low** |
| Grassland - shrub | 141 | 154 | 84 |
| Agriculture field, long-term fallow | 18 | 15 | 21 |
| Mixed<sup>b</sup> | 22 | 9 | 29 |
| Juniper woodland - grass | 0 | 6 | 30 |
| Pine forest | 6 | 6 | 9 |
| Pinyon juniper woodland - grass | 6 | 0 | 18 |
| Grassland - juniper | 8 | 0 | 0 |
| Agriculture field, active | 29 | 0 | 0 |

<sup>a</sup>For the hyphenated ecosystem types, the dominant plant type is listed first and the minor plant type is listed after the hyphen.
<sup>a</sup>"Mixed" represented locations that were predominantly low-lying drainage areas (e.g. arroyos, vales) with highly mixed plant types (trees, shrubs, and grasses).

### Relationship between community-level features and DOC abundance

The composition of both the fungal and bacterial communities in the low DOC microcosms at the end of the 44-day incubation was significantly different from the composition of those in the high DOC microcosms (PERMANOVA:bacteria, F_1,309_ = 14.48, *P* = <0.001; fungi, F_1,343_ = 7.64, *P* = <0.001; Fig. 3, Suppl. Fig. 4). The mantel correlation between DOC abundance and community composition was stronger for bacteria (Mantel test; r = 0.38, *P* = <0.001) than for fungi (Mantel test; r = 0.12, *P* = <0.001). On average, 23 bacterial orders containing a minimum of 0.1% of the bacterial sequences were observed per sample, and 57% of these differed significantly in relative abundances between the low and high DOC groups (Suppl. Fig. 4A). There were 23 bacterial families containing at least 1% of the sequences on average, and 13 of these were significantly different in relative abundances between the two groups (Suppl. Fig. 4B). For fungi, 14 orders contained at least 0.1% of the fungal sequences per sample, and only 36% of these differed significantly between the low and high DOC groups, further reflecting the stronger link between bacterial community composition and DOC levels (Suppl. Fig. 4C).

**Fig. 3.**
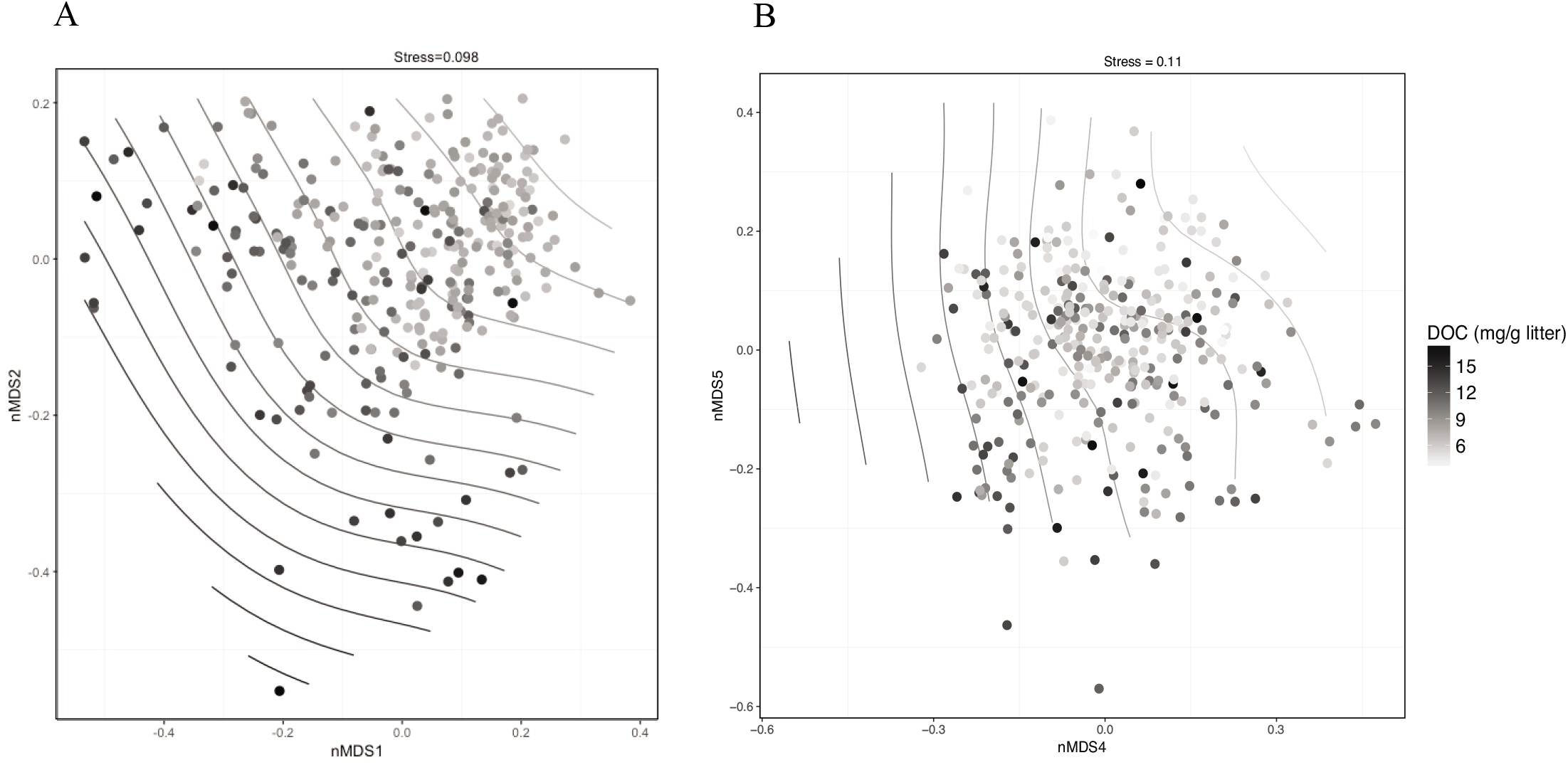
Relationship between microbial community composition and dissolved organic carbon (DOC) abundance. Non-metric multidimensional scaling ordinations performed on rarefied data for (A) bacterial communities (dimensions one and two, 1023 sequences per sample) and (B) fungal communities (dimensions four and five, 2032 sequences per sample) overlain on a smoothed surface showing variation in DOC quantity. Microbial communities developed in microcosms during 44 days of pine litter decomposition. Points are shaded by DOC abundance. The stress value is derived from six dimensions.

Further community-level analyses were undertaken using DNA extracted from the high and low DOC microcosms. High DOC microcosm communities had, on average, 18% less biomass (measured as total extracted DNA) than low DOC communities (two-tailed *t*-test, *t*_378_ = 4.7, *P* = <0.001; Suppl. Fig.5) but DOC was only weakly correlated with biomass (Pearson correlation; r = −0.22, *P* = <0.001). The ratio of fungi to bacteria estimated from total DNA and qPCR data from a subset of samples was not significantly different between high and low DOC communities (two-tailed *t*-test, *t*_36_ = 1.7, *P* = 0.1). The community-level trait most strongly linked to DOC abundance in our study was bacterial richness (Pearson correlation; r = −0.64, *P* = <0.001). The average bacterial richness was one third lower in high DOC communities compared to low DOC communities (Fig. 4A, two-tailed *t* test, *t*_307_ = 13.74, *P* = <0.001), whereas fungal richness was not significantly different (Fig. 4b). Comparisons of diversity between high and low groups made using the Shannon-Weiner index yielded similar results (bacteria: two-tailed *t* test, t_288_ = 10.39, *P* = <0.001, fungi: two-tailed *t* test, t_331_ = 0.24, *P* = 0.81).

**Fig. 4.**
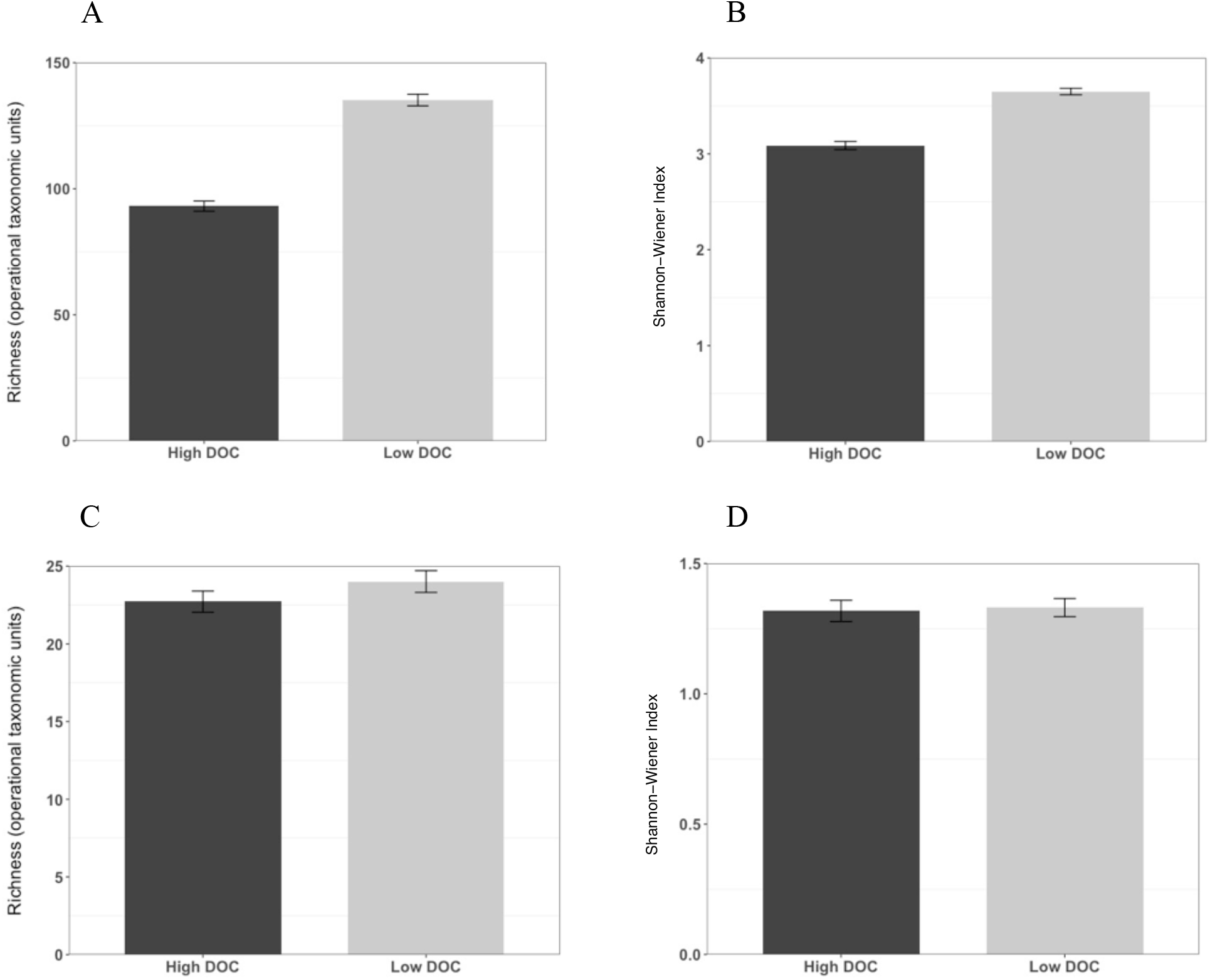
Operational taxonomic unit (OTU) richness for bacteria (A) and fungi (C), and Shannon-Wiener diversity for bacteria (B) and fungi (D), in high and low DOC microcosms. For bacteria, *n* = 311 and for fungi, *n* = 345. Communities developed during 44 days of pine litter decomposition. The data are expressed as the mean value ± SEM. The differences in bacterial richness and Shannon-Weiner values were significant, as determined by two-tailed *t* tests (*P* = <0.001).

## Discussion

Laboratory microcosms lack many elements of natural systems. Nonetheless, microcosms are a useful testbed to discover basic phenomena that merit further investigation in natural systems. Although further research is required to determine the true relevance of our microcosm results for natural ecosystems, it is useful to consider the potential implications, which is our focus in the following sections.

### Variation in carbon flows from decomposition

By holding the environment constant within laboratory microcosms and varying nothing but microbial community composition, we demonstrated an incontrovertible link between microbial community composition and decomposition outcomes. Our findings build upon those of Strickland et al. (27) and Matulich & Martiny (28), being generated from much larger numbers of naturally assembled microbial communities. In addition, we found the range of variation in CO_2_ and DOC outputs is substantial enough to merit further study in relation to climate feedbacks. This range, combined with the general magnitude of natural CO_2_ flux from soil respiration (29), suggests the possibility of exploiting microbial-driven variation in CO_2_ output from soils to achieve the proposed offsets in anthropogenic CO_2_ emissions needed to mitigate climate change (30). We acknowledge our current results merely illustrate a conceptual possibility. The true range in CO_2_ flux that can arise from natural or manipulated microbial community variation within a natural ecosystem remains unknown. Given that surface litter decomposition is only one component of the carbon cycle in soil, a wide range of microbial-driven variation in surface litter decomposition may routinely be counter-balanced in nature by compensatory processes over longer time-scales (31) or in other components of the carbon cycle, such that an ecosystem will exhibit a fairly stable mean CO_2_ flux. Nonetheless, our findings illustrate an important phenomenon that motivates further investigation of the naturally occurring range in microbial-driven CO_2_ flux, as well as the impact of maintaining specific patterns of carbon flow over long time scales by manipulating microbial community composition.

We focused on the DOC distribution for the determination of community traits important to carbon flows in soil. The DOC distribution we detected suggests a large potential for microbial community control over soil carbon storage because DOC has the possibility for transport from the soil surface to deep soil horizons where long-term carbon storage can occur (32). When DOC from decomposing surface litter is transported to deeper layers, some of the carbon adsorbs to mineral surfaces (33, 34) enabling carbon storage over millennial timescales (35, 36). Consequently, in natural systems, DOC from surface litter contributes significantly to soil carbon stocks (37). Because the amount of carbon stored is related to the quantity of DOC available for translocation (37), microbial communities that yield larger quantities of DOC create a possibility for greater soil carbon storage.

In addition to changing DOC quantity, our results show that microbial community composition can alter DOC quality. The finding that communities with high quantities of DOC also had, on average, DOC with higher mineral binding potential suggests further possibilities for impacts of microbial community composition on DOC storage in soil. Increases in the mineral-binding fraction of DOC involve enrichment of DOC with compounds that have affinity for mineral surfaces (32). Enrichment of DOC may occur through different mechanisms including (a) variable depletion of compounds released from plant litter, (b) production of taxon-specific microbial by-products (e.g. polyphenolics produced by Actinobacteria) (38) and (c) release of taxon-specific residues from dead microbial cells such as melanin, chitin, B-glucans, or glycoproteins (e.g. glomalin) from fungi (39–41). Combining the effects of DOC quantity and quality, we observed a 7-fold range in readily-storable carbon among the microbial communities in our study system. In a natural system, the *realized* quantity of carbon stored would depend on additional factors such as precipitation enabling DOC transport to deep mineral layers (42), soil porosity (43), the minerology and chemistry of soil (44), and variation in the composition of subsurface microbial communities that control the extent of DOC decomposition during DOC transport through the soil (45). Given the complexity and heterogeneity of natural systems, the range of microbially-driven variation in storable carbon could be either smaller or larger than the range observed in our system, and further study in natural systems is needed to determine this.

The geographic intermingling of the source soils that produced communities yielding extreme (high or low) DOC quantities in our microcosm study suggests natural communities may vary enough over small spatial scales (meters to kilometers) to create substantial variation in decomposition outcomes and, therefore, climate feedbacks. The entire functional distribution we observed with DOC may occur within single ecosystems in nature, facilitated by patchiness in microbial community composition at the centimeter to meter scale (46, 47). Up to 70-fold variation in carbon-mass loss during litter decomposition has been recorded within single grassland sites (48), and could conceivably have arisen from variation in the composition of decomposer communities. These observations emphasize the need for explicit analysis of microbial composition as a driver when substantial variation in carbon flow is seen in a natural ecosystem. If wide a wide range of compositional and functional variation in surface decomposer communities routinely occurs within ecosystems, it would suggest the absence of strongly deterministic processes that would force functional convergence. Confirming such plasticity would increase the possibility of successfully steering communities to achieve desired patterns of carbon flow.

### Community-level features linked to carbon flow

To facilitate identification of community-level features linked to DOC abundance, we sequenced the subset of microbial communities that represented the tails of the DOC abundance distribution, delineated as contrasting functional states (high and low DOC quantity). We focused on four community-level features that are often measured in ecosystem studies: 1) community biomass, 2) the estimated ratio of fungi to bacteria, 3) fungal and bacterial community composition, and 4) fungal and bacterial richness and diversity.

We found that high DOC communities from microcosms at the end of the 45-day incubation had less biomass (measured as total extracted DNA) than low DOC communities, but DOC was not strongly correlated with biomass. The small range of variation we observed in microbial biomass could conceivably arise from differences in community composition, given the large (e.g. >2-fold) variation in genome size that occurs among bacterial species and among fungal species. The ratio of fungi to bacteria was not significantly different between groups although this trait has been reported as an indicator of soil carbon content (49). However significant differences were found both in fungal and bacterial composition between the high versus low DOC communities. Fungi are generally considered the main microbial drivers of plant litter decomposition, due to their production of powerful enzymes for deconstruction of plant lignocellulose (50). In our study, however, the correlation between DOC abundance and community composition was weaker for fungi than for bacteria. Bacterial communities are also important to decomposition outcomes in the field (31). Our results are consistent with the view that fungi drive the overall rate and extent of litter deconstruction, while bacteria play a significant role as secondary consumers, controlling the quantity of DOC that remains available for transport into soil.

In addition to the importance of bacterial community composition to DOC levels, we found a strong link between DOC and bacterial richness and diversity. Fungal richness and diversity were not found to be important. The correspondence of lower bacterial richness with higher DOC abundance may indicate either an absence of critical functions (e.g. genes required for the consumption of specific DOC compounds or concentrations) in the microbial communities, or suppression of those critical functions. Functional suppression can arise from negative interactions within microbial communities, such as antibiotic production (51), predation (52), or bacteriophage activity (53). Bacterial richness is known to vary at the landscape scale, declining with greater aridity in at least some biomes (54, 55), and with lower pH (56). If bacterial richness proves to be a robust factor in DOC abundance among natural ecosystems, understanding the interplay between bacterial richness, taxonomic composition, and carbon flow may reveal mechanisms that can be integrated in models, improving prediction of soil carbon stocks across a variety of environments.

### Summary

To improve climate predictions by including microbial processes in soil carbon models, climate-relevant microbial processes and simple traits that represent them must first be identified, as has been achieved with plant traits (57, 58). Our study showed a strong influence of microbial community composition over decomposition outcomes in a constant environment, resulting in large differences in carbon flow from litter decomposition. It is reasonable to expect that microbial composition drives variation in every component of soil carbon cycling (e.g. surface litter decomposition, subsurface litter decomposition, plant productivity and carbon allocation). Our findings motivate investigation of this phenomenon in natural systems in order to assess its importance to climate feedbacks within and among existing ecosystems and its implications for managing soil carbon. We identified a high-level trait, bacterial richness, linked to DOC abundance and known to be geographically patterned. Bacterial richness has also been linked to carbon fate in mammals (59) where lower richness correlates with increased carbon storage in the host (59, 60). Our findings raise the tantalizing possibility of discovering robust principles that underpin functional states in extremely diverse systems ranging from soils to animal guts.

## Acknowledgements

This work was supported by grant SFA 2015LANLF260 from the US Department of Energy Office of Biological and Environmental Research.

## References

1. Bradford MA, Berg B, Maynard DS, Wieder WR, Wood SA (2016) Understanding the dominant controls on litter decomposition. J Ecol 104: 229–238

2. Melillo JM, Frey SD, DeAngelis KM, Werner WJ, Bernard MJ, Bowles FP, Pold G, Knorr MA, Grandy AS (2017) Long-term pattern and magnitude of soil carbon feedback to the climate system in a warming world. Science 358: 101–105

3. van der Wal A, Geydan TD, Kuyper TW, de Boer W (2013) A thready affair: linking fungal diversity and community dynamics to terrestrial decomposition processes. FEMS Microbiol Rev 37: 477–494

4. Bier RL, Bernhardt ES, Boot CM, Graham EB, Hall EK, Lennon JT, Nemergut DR, Osborne BB, Ruiz-Gonzalez C, Schimel J, Waldrop MP, Wallenstein MD (2015) Linking microbial community structure and microbial processes: an empirical and conceptual overview. FEMS Microbiol Ecol 91: fiv113

5. Blasche S, Kim Y, Oliveira AP, Patil KR (2017) Model microbial communities for ecosystems biology. Curr Opin Syst Biol 6: 51–57

6. Jessup CM, Kassen R, Forde SE, Kerr B, Buckling A, Rainey PB, Bohannan BJM (2004) Big questions, small worlds: microbial model systems in ecology. Trends Ecol Evol 19: 189–197

7. Powell JR, Karunaratne S, Campbell CD, Yao H, Robinson L, Singh BK (2015) Deterministic processes vary during community assembly for ecologically dissimilar taxa. Nat Commun 6: 8444

8. Cotrufo MF, Soong JL, Horton AJ, Campbell EE, Haddix ML, Wall DH, Parton WJ (2015) Formation of soil organic matter via biochemical and physical pathways of litter mass loss. Nature Geosci 8: 776–779

9. Müller K, Marhan S, Kandeler E, Poll C (2017) Carbon flow from litter through soil microorganisms: From incorporation rates to mean residence times in bacteria and fungi. Soil Biol Biochem 115: 187–196

10. Bates ST, Berg-Lyons D, Caporaso JG, Walters WA, Knight R, Fierer N (2010) Examining the global distribution of dominant archaeal populations in soil. Isme J 5: 908

11. Mueller RC, Gallegos-Graves LV, Kuske CR (2016) A new fungal large subunit ribosomal RNA primer for high-throughput sequencing surveys. FEMS Microbiol Ecol 92: e153

12. Talbot JM, Bruns TD, Taylor JW, Smith DP, Branco S, Glassman SI, Erlandson S, Vilgalys R, Liao H-L, Smith ME, Peay KG (2014) Endemism and functional convergence across the North American soil mycobiome. PNAS 111: 6341–6346

13. Mueller R, Belnap J, Kuske C (2015) Soil bacterial and fungal community responses to nitrogen addition across soil depth and microhabitat in an arid shrubland. Front Microbiol 6: e891

14. Gloor GB, Hummelen R, Macklaim JM, Dickson RJ, Fernandes AD, MacPhee R, Reid G. (2010) Microbiome profiling by Illumina sequencing of combinatorial sequence-tagged PCR products. PLoS ONE 5: e15406

15. Zhang J, Kobert K, Flouri T, Stamatakis A (2014) PEAR: a fast and accurate Illumina Paired-End reAd mergeR. Bioinformatics 30: 614–20

16. Caporaso JG, Kuczynski J, Stombaugh J, Bittinger K, Bushman FD, Costello EK, et al (2010) QIIME allows analysis of high-throughput community sequencing data. Nat Methods 7: 335

17. Edgar RC (2013) UPARSE: highly accurate OTU sequences from microbial amplicon reads. Nat Methods 10: 996

18. Edgar RC, Haas BJ, Clemente JC, Quince C, Knight R (2011) UCHIME improves sensitivity and speed of chimera detection. Bioinformatics 27: 2194–2200

19. Wang Q, Garrity GM, Tiedje JM, Cole JR (2007) Naive bayesian classifier for rapid assignment of rRNA sequences into the new bacterial taxonomy. Appl Environ Microbiol 73: 5261–5267

20. Lane DJ (1991) 16S/23S rRNA sequencing. In: Stackebrandt E, Goodfellow M (ed), Nucleic acid techniques in bacterial systematics. John Wiley & Sons, Chichester, UK

21. Muyzer G, de Waal EC, Uitterlinden AG (1993) Profiling of complex microbial populations by denaturing gradient gel electrophoresis analysis of polymerase chain reaction-amplified genes coding for 16S rRNA. Appl Environ Microbiol 59: 695–700

22. Castro HF, Classen AT, Austin EE, Norby RJ, Schadt CW (2010) Soil microbial community responses to multiple experimental climate change drivers. Appl Environ Microbiol 84: 999–1007

23. Oksanen J, Blanchet FG, Friendly M, Kindt R, Legendre R, McGlinn D (2017) vegan: Community Ecology Package. R package version 2.4-3

24. Anderson MJ (2001) A new method for non-parametric multivariate analysis of variance. Austral Ecology 26: 32–46

25. Goslee SC, Urban DL (2007) The ecodist package for dissimilarity-based analysis of ecological data. J Stat Softw 22: 1–19

26. R Core Team (2017) R: A language and environment for statistical computing, R Foundation for Statistical Computing, Vienna, Austria. https://www.R-project.org/

27. Strickland MS, Lauber C, Fierer N, Bradford MA (2009) Testing the functional significance of microbial community composition. Ecology 90: 441–451

28. Matulich KL, Martiny JB (2015) Microbial composition alters the response of litter decomposition to environmental change. Ecology 96: 154–163

29. Raich JW, Schlesinger WH (1992) The global carbon dioxide flux in soil respiration and its relationship to vegetation and climate. Tellus B 44: 81–99

30. Gao Y, Gao X, Zhang X (2017) The 2 °C global temperature target and the evolution of the long-term goal of addressing climate change—from the United Nations Framework Convention on Climate Change to the Paris Agreement. Engineering 3: 272–278

31. Glassman SI, Weihe C, Li J, Albright MBN, Looby CI, Martiny AC, Treseder KK, Allison SD, Martiny JBH (2018) Decomposition responses to climate depend on microbial community composition. PNAS 115: 11994–11999

32. Kleber M, Eusterhues K, Keiluweit M, Mikutta C, Mikutta R, Nico PS (2015) Mineral-Organic Associations: Formation, properties, and relevance in soil environments. Adv Agron 130: 1–140

33. Newcomb CJ, Qafoku NP, Grate JW, Bailey VL, De Yoreo JJ (2017) Developing a molecular picture of soil organic matter–mineral interactions by quantifying organo–mineral binding. Nat Commun 8: 396

34. Kaiser K, Guggenberger G (2000) The role of DOM sorption to mineral surfaces in the preservation of organic matter in soils. Org Geochem 31: 711–725

35. Rumpel C, Kögel-Knabner I, Bruhn F (2002) Vertical distribution, age, and chemical composition of organic carbon in two forest soils of different pedogenesis. Org Geochem 33: 1131–1142

36. Schöning I, Kögel-Knabner I. 2006. Chemical composition of young and old carbon pools throughout Cambisol and Luvisol profiles under forests. Soil Biol Biochem 38: 2411–2424

37. Kalbitz K, Kaiser K (2008) Contribution of dissolved organic matter to carbon storage in forest mineral soils. J Plant Nutr Soil Sc 171: 52–60

38. Trigo C, Ball AS. 1994. Is the solubilized product from the degradation of lignocellulose by actinomycetes a precursor of humic substances? Microbiology 140: 3145–3152

39. Kögel-Knabner I (2002) The macromolecular organic composition of plant and microbial residues as inputs to soil organic matter. Soil Biol Biochem 34: 139–162

40. Fernandez CW, Koide RT (2012) The role of chitin in the decomposition of ectomycorrhizal fungal litter. Ecology 93: 24–28

41. Siletti CE, Zeiner CA, Bhatnagar JM (2017) Distributions of fungal melanin across species and soils. Soil Biol Biochem 113: 285–293

42. Neff JC, Asner GP (2001) Dissolved organic carbon in terrestrial ecosystems: synthesis and a model. Ecosystems 4: 29–48

43. Bailey VL, Smith AP, Tfaily M, Fansler SJ, Bond-Lamberty B (2017) Differences in soluble organic carbon chemistry in pore waters sampled from different pore size domains. Soil Biol Biochem 107: 133–143

44. Doetterl S, Stevens A, Six J, Merckx R, Van Oost K, Casanova Pinto M, Casanova-Katny A, Muñoz C, Boudin M, Zagal Venegas E, Boeckx P (2015) Soil carbon storage controlled by interactions between geochemistry and climate. Nat Geosci 8: 780

45. Dong W, Wan J, Tokunaga TK, Gilbert B, Williams KH (2017) Transport and humification of dissolved organic matter within a semi-arid floodplain. J Environ Sci 57: 24–32

46. Štursová M, Bárta J, Šantrůčková H, Baldrian P (2016) Small-scale spatial heterogeneity of ecosystem properties, microbial community composition and microbial activities in a temperate mountain forest soil. FEMS Microbiol Ecol 92: fiw185–fiw185

47. O’Brien SL, Gibbons SM, Owens SM, Hampton-Marcell J, Johnston ER, Jastrow JD, Gilbert JA, Meyer F, Antonopoulos DA (2016) Spatial scale drives patterns in soil bacterial diversity. Environ Microbiol 18: 2039–2051

48. Bradford MA, Veen GF, Bonis A, Bradford EM, Classen AT, Cornelissen JHC, et al (2017) A test of the hierarchical model of litter decomposition. Nat Ecol Evol 1: 1836–1845

49. Malik AA, Chowdhury S, Schlager V, Oliver A, Puissant J, Vazquez PGM, Jehmlich N, von Bergen M, Griffiths RI, Gleixner G (2016) Soil fungal:bacterial ratios are linked to altered carbon cycling. Front Microbiol 7:e1247

50. Baldrian P (2017) Forest microbiome: diversity, complexity and dynamics. FEMS Microbiol Rev 41: 109–130

51. Frey-Klett P, Burlinson P, Deveau A, Barret M, Tarkka M, Sarniguet A (2011) Bacteria-Fungal interactions: Hyphens between agricultural, clinical, environmental and food microbiologists. Microbiol Mol Biol Rev 75: 583–609

52. Sockett RE. (2009) Predatory lifestyle of Bdellovibrio bacteriovorus. Annu Rev Microbiol 63: 523–539

53. Williamson KE, Fuhrmann JJ, Wommack KE, Radosevich M (2017) Viruses in soil ecosystems: an unknown quantity within an unexplored territory. Annu Rev Virol 4: 201–219

54. Tu B, Domene X, Yao M, Li C, Zhang S, Kou Y, Wang Y, Li X (2017) Microbial diversity in Chinese temperate steppe: unveiling the most influential environmental drivers. FEMS Microbiol Ecol 93: e031

55. Maestre FT, Delgado-Baquerizo M, Jeffries TC, Eldridge DJ, Ochoa V, Gozalo B, et al (2015) Increasing aridity reduces soil microbial diversity and abundance in global drylands. PNAS 112: 15684–9

56. Bahram M, Hildebrand F, Forslund SK, Anderson JL, Soudzilovskaia NA, Bodegom PM, et al (2018) Structure and function of the global topsoil microbiome. Nature 560: 233–237

57. Laughlin DC. 2014. The intrinsic dimensionality of plant traits and its relevance to community assembly. J Ecol 102: 186–193

58. Wieder WR, Grandy AS, Kallenbach CM, Taylor PG, Bonan GB (2015) Representing life in the Earth system with soil microbial functional traits in the MIMICS model. Geosci Model Dev 8: 1789–1808

59. Shabat SKB, Sasson G, Doron-Faigenboim A, Durman T, Yaacoby S, Berg Miller ME, White BA, Shterzer N, Mizrahi I (2016) Specific microbiome-dependent mechanisms underlie the energy harvest efficiency of ruminants. ISME J 10: 2958–2972

60. Le Chatelier E, Nielsen T, Qin J, Prifti E, Hildebrand F, Falony G, Almeida M, Arumugam M, Batto J-M, Kennedy S (2013) Richness of human gut microbiome correlates with metabolic markers. Nature 500: 541–546

